# Pushed to extremes: distinct effects of high temperature vs. pressure on the structure of an atypical phosphatase

**DOI:** 10.1101/2023.05.02.538097

**Authors:** Liliana Guerrero, Ali Ebrahim, Blake T. Riley, Minyoung Kim, Qingqiu Huang, Aaron D. Finke, Daniel A. Keedy

**Author notes:** contributed equally (Liliana Guerrero, Ali Ebrahim).

## Abstract

Protein function hinges on small shifts of three-dimensional structure. Elevating temperature or pressure may provide experimentally accessible insights into such shifts, but the effects of these distinct perturbations on protein structures have not been compared in atomic detail. To quantitatively explore these two axes, we report the first pair of structures at physiological temperature vs. high pressure for the same protein, STEP (PTPN5). We show that these perturbations have distinct and surprising effects on protein volume, patterns of ordered solvent, and local backbone and side-chain conformations. This includes novel interactions between key catalytic loops only at physiological temperature, and a distinct conformational ensemble for another active-site loop only at high pressure. Strikingly, in torsional space, physiological temperature shifts STEP toward previously reported active-like states, while high pressure shifts it toward a previously uncharted region. Together, our work argues that temperature and pressure are complementary, powerful, fundamental macromolecular perturbations.

## Introduction

The biological functions of many proteins require transitions between conformational substates ^1–3^. Despite their functional importance, protein conformational substates are often difficult to characterize. X-ray crystallography can prove useful in this regard by revealing alternate conformations that coexist in crystals at partial occupancy, as shown by electron density maps ^4^. Such conformational ensembles can be shifted by discrete, localized, targeted perturbations like ligands ^5^ or mutations ^6^, revealing insights into allostery and enzyme catalysis. However, known ligands are unavailable for most sites in most proteins, and predicting the effects of mutations is difficult. By contrast, continuous, global, generic biophysical perturbations offer advantages: they can be applied to any protein, affect the entire structure simultaneously, and can be “titrated” to shift conformational distributions and map correlated conformational changes relevant to function ^7^.

One such biophysical perturbation, which has gained traction as a valuable experimental variable in structural biology and biophysics, is temperature (T). Room-temperature (RT) X-ray crystallography ^8^ avoids structural biases of cryogenic-temperature crystallography, revealing differences in protein conformation ^9–12^, ligand binding ^13,14^, and solvation layers ^10,14^. Multitemperature crystallography provides additional insights into conformational coupling ^11,12,15^. Notably, crystal structures at physiological temperature (37°C, 310 K) can reveal unique protein conformations ^12,16^. RT crystallography methods are rapidly improving ^8^, including serial crystallography ^17^. RT crystal structures are also increasingly used in computational simulations ^18–20^.

Complementary to temperature, but relatively underexplored, is pressure (P). Whereas high temperature stabilizes states with high entropy, high pressure stabilizes states with low volume, isolating distinct excited states that may have unique links to biological function ^21–25^. Importantly, pressure-induced structural changes on the sub-angstrom level observed by high-pressure X-ray crystallography have been shown to be directly related to protein function ^26^. Other past high-pressure protein crystallography studies showed non-uniform responses of coordinates and B-factors ^27^, non-compressive conformational shifts mirroring those induced by pH change ^28^, water infiltration into engineered ^29,30^ and natural cavities ^31^, crystal phase transitions ^31,32^, conformational shifts of functional residues in an allosteric network ^32^, and changes in ligand affinity ^33^.

Despite this foundation, relatively few studies have explored the detailed effects of pressure on protein conformational ensembles using crystallography. While a few studies have highlighted protein alternate conformations for isolated residues ^31,32^, to our knowledge no study has comprehensively explored the effects of pressure on detailed conformational ensembles of all residues throughout a protein structure. Moreover, very few studies ^28^ have compared the atomic-level effects of elevated temperature vs. pressure on protein crystal structures. It thus remains unclear whether, and how, these two fundamental thermodynamic perturbations differentially affect protein conformational ensembles, which limits our toolkit for probing fundamental connections between conformational heterogeneity and biological function.

An attractive system to investigate the differential effects of temperature vs. pressure on protein conformational ensembles is the protein tyrosine phosphatase (PTP) enzyme STEP (PTPN5). STEP is a brain-specific PTP, and a validated therapeutic target for Alzheimer’s disease ^34^, Fragile X syndrome ^35^, and Parkinson’s disease ^36^. The public Protein Data Bank ^37^ includes 7 high-resolution (1.66-2.15 Å) crystal structures of human STEP with different ligands, demonstrating its tractability with crystallography. As revealed in these structures, STEP has several unusual features among PTPs, including an “atypically open” active-site WPD loop conformation ^38^ and an allosteric site with a small-molecule activator (not inhibitor) ^39^. However, all existing STEP structures are at cryogenic temperature and ambient pressure.

Here we report high-resolution (< 2 Å) crystal structures of unliganded STEP at high temperature (HiT) and at high pressure (HiP), along with a reference structure at low temperature and low pressure (LoTP). To our knowledge, these new structures of STEP represent several firsts. Our high-temperature structure is only the eleventh crystal structure of any protein, and the first of any phosphatase (or kinase), at physiological temperature or above (≥ 310 K). Our high-pressure structure of STEP is also the first of any phosphatase (or kinase) at high pressure. Together, our new structures make STEP the first protein with crystal structures at both physiological temperature and high pressure, presenting a unique opportunity to compare the effects of these two distinct perturbations on protein conformational ensembles.

By quantitatively interrogating these data, we reveal that temperature and pressure have complementary effects on the conformational landscape of STEP. These two perturbations have opposite effects on the crystal lattice but surprisingly similar effects on the protein molecular volume, stabilize distinct ordered water molecules throughout the protein, induce backbone shifts in non-overlapping regions of the structure, and rearrange different sets of side chains. We observe a previously unseen arrangement of product-like anions in the active-site pocket, new conformations of conserved catalytic residues only at high temperature, and an active-like conformation of an active-site loop only at high pressure. Surprisingly, using a new computational method for analyzing distributions of protein structures ^40^, we find that high temperature in the apo state induces a coordinated global shift toward previous ligand-bound active-like structures, whereas high pressure shifts the protein toward a previously unseen region of conformational space. Overall, our results illustrate the potential of manipulating protein structures with a broad spectrum of physical perturbations to gain unique insights into their mechanical coupling and biological function.

## Results

### Overview of STEP structure and active site

As a reference point for comparing the effects of temperature vs. pressure on STEP, we first determined a traditional crystal structure of the STEP catalytic domain at cryogenic temperature (100 K) and ambient pressure (0.1 MPa). The diffraction dataset and resulting refined structure were of high quality (**Table 1**). This structure has the expected PTP catalytic domain architecture (**Fig. 1**), including several key loops surrounding the active site region (**Fig. 1c**). Broadly speaking, it is similar to the 7 previously published structures of human STEP (8 including 1 structure of mouse STEP) (**Supp. Fig. 1**).

**Table 1:** Crystallographic statistics. Overall statistics given first (statistics for highest-resolution bin given in parentheses).

| PDB ID | 8SLS | 8SLT | 8SLU |
| --- | --- | --- | --- |
| Dataset name | LoTP | HiT | HiP |
| Temperature (K) | 100 | 310 | 100 |
| Pressure (MPa) | 0.1 (ambient) | 0.1 (ambient) | 205 |
| Resolution (Å) | 1.71-67.58 | 1.96-68.61 | 1.84-57.37 |
| Completeness (%) | 98.64 (93.42) | 99.80 (99.03) | 99.45 (99.12) |
| Multiplicity | 4.7 (4.8) | 6.7 (6.9) | 6.6 (6.6) |
| I/sigma(I) | 10.23 (0.59) | 7.79 (0.63) | 10.71 (0.64) |
| R<sub>merge</sub>(I) | 0.108 (2.37) | 0.170 (2.398) | 0.114 (2.127) |
| R<sub>meas</sub>(I) | 0.122 (2.62) | 0.184 (2.593) | 0.124 (2.308) |
| R<sub>pim</sub>(I) | 0.055 (1.106) | 0.071 (0.976) | 0.049 (0.888) |
| CC<sub>1/2</sub> | 0.96 (0.374) | 0.996 (0.358) | 0.998 (0.687) |
| Total observations | 176675 (17614) | 176185 (17871) | 196146 (19601) |
| Unique observations | 37552 (3655) | 26353 (2573) | 29880 (2949) |
| Space group | P212121 | P212121 | P212121 |
| Unit cell dimensions (Å, Å, Å, °, °, °) | 39.67, 63.51, 135.16, 90, 90, 90 | 39.98, 64.49, 137.21, 90, 90, 90 | 39.14, 63.47, 134.20, 90, 90, 90 |
| Solvent content (%) | 48.68 | 50.60 | 47.58 |
| R<sub>work</sub> (%) | 19.08 | 17.34 | 19.88 |
| R<sub>free</sub> (%) | 22.02 | 21.12 | 23.33 |
| RMS bonds (Å) | 0.016 | 0.012 | 0.011 |
| RMS angles (°) | 1.24 | 1.062 | 1.103 |
| Ramachandran outliers (%) | 0.36 | 0.36 | 0.36 |
| Ramachandran favored (%) | 94.64 | 95.36 | 94.29 |
| Clashscore | 2.08 | 1.95 | 5.64 |
| MolProbity score | 1.44 | 1.28 | 1.69 |

**Figure 1:**
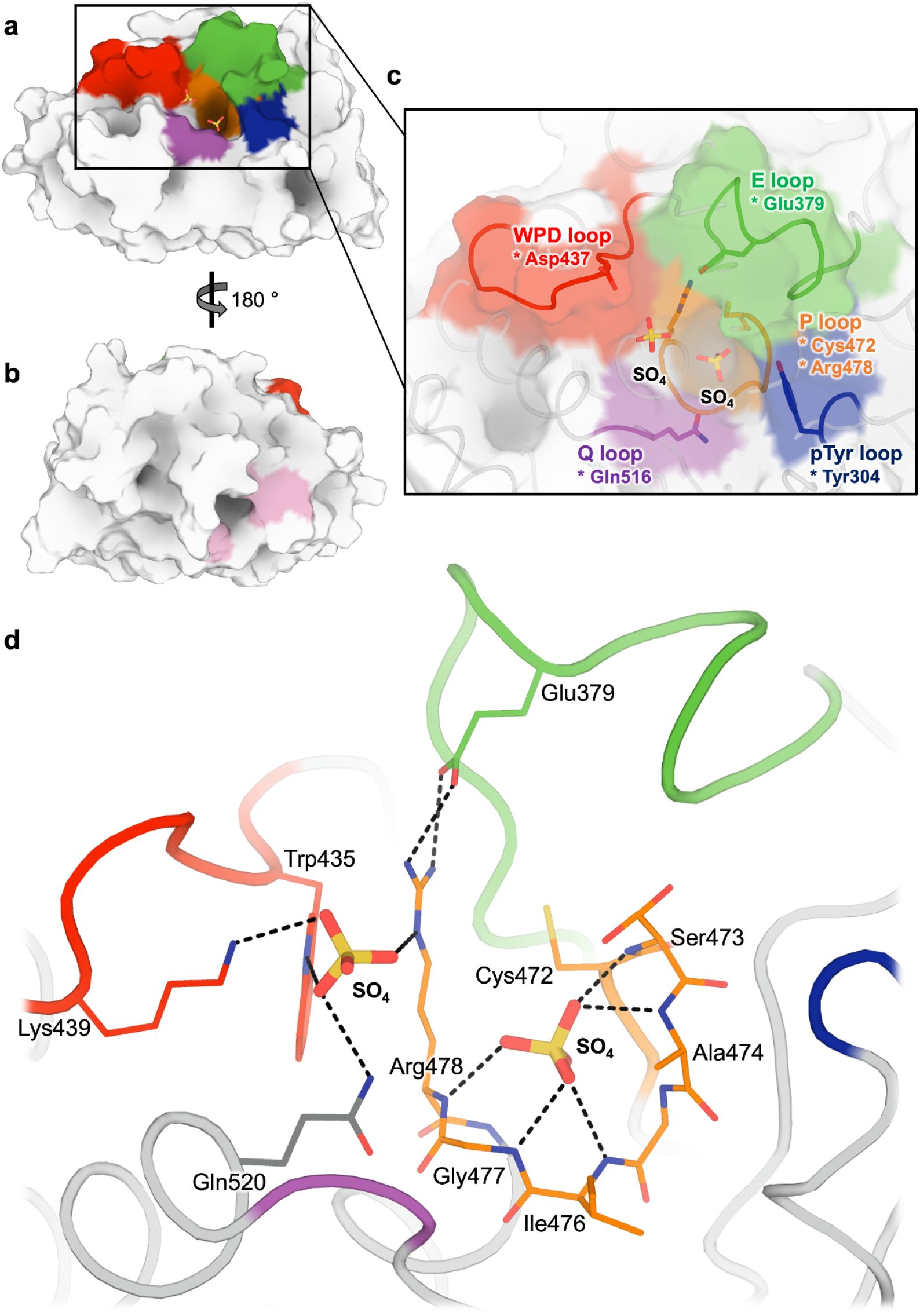
Structural overview of STEP, including two sulfates in active site. a) Overview of STEP catalytic domain, centered on active site. b) 180° rotation of (a) to show allosteric activator binding site ^39^, with key residues highlighted in pink. c) Zoom-in of (a) showing several key active-site loops and two sulfates bound in the active site cleft. Key catalytic residues are denoted with an asterisk. d) Interactions between two sulfates and nearby residues in the active site of our LoTP structure.

One notable feature of this new structure (as well as our high-temperature and high-pressure structures, introduced below) differs from the previous STEP structures: the active site binds two sulfate molecules (**Fig. 1c-d, Supp. Fig. 2a**). The “bottom” sulfate is well-coordinated by the catalytic P loop, analogous to the phosphate group in the phosphotyrosine (pTyr) substrate in PDB ID 2cjz ^38^ (**Supp. Fig. 2c**). This sulfate positioning is further supported by a new side-chain rotamer for the catalytic Cys472 to avoid a steric clash to the sulfate, which is present in our new structures but previously was only seen as a partial-occupancy alternate conformation in an allosterically activated structure (PDB ID 6h8r) (**Supp. Fig. 2b**). The “top” sulfate sits just beneath the catalytic WPD loop, where a lone sulfate has been observed previously in PDB ID 2bv5, 2bij ^41^, and 6h8r ^39^, and inhibitors with negatively charged moieties have been observed in PDB ID 5ovr, 5ovx, and 5ow1 ^42^ (**Supp. Fig. 2b**). Thus, although previous structures of STEP have sulfates or phosphate-like chemical groups independently in each of these sites, no previous structure has them in both sites simultaneously. The closest comparison is PDB ID 2bv5, in which the catalytic Cys472 is modeled as acetylated in the bottom site and a sulfate is in the top site (**Supp. Fig. 2d**), but this arrangement differs in chemical character from what we observe.

### X-ray datasets at high temperature vs. pressure

To complement our reference structure of STEP at low temperature and low pressure, we used similarly prepared crystals to obtain two additional crystal structures: one at physiological temperature (310 K) but ambient pressure, and another at high pressure (205 MPa) but cryogenic temperature via high-pressure cryocooling ^43^. For the remainder of this paper, we refer to the structure at low temperature and low pressure as LoTP, the structure at high temperature as HiT, and the structure at high pressure as HiP. The HiT and HiP diffraction datasets and resulting refined structures were of similarly high quality as for LoTP, including acceptably similar resolutions (**Table 1**). Therefore, these datasets can be directly compared to gain insights into the differential effects of temperature vs. pressure on the conformational ensemble of STEP.

The unit cell dimensions differed from the LoTP reference dataset in opposing ways: at HiT the unit cell volume expanded by 3.9%, whereas at HiP the unit cell was instead compressed by 2.1% (**Table 2**). Comparatively, the protein molecule itself was more robust, but was still affected by temperature and pressure: at HiT the protein volume expanded by 1.2%, whereas at HiP it still expanded, but by only 0.5%. Thus elevated temperature expands the unit cell and, to a lesser extent, the protein itself; by contrast, elevated pressure compresses the unit cell, but still allows the protein itself to slightly expand. These observations suggest that temperature vs. pressure have more complex effects on STEP than might be naïvely expected from the unit cell changes alone. Indeed, these distinct perturbations induce a variety of conformational changes distributed throughout the structure of STEP, including some with potential biological relevance, as shown below.

**Table 2:** Change in unit cell and protein volume at high temperature vs. high pressure. Absolute number given first (% change relative to LoTP given in parentheses). Protein total volume calculated by the ProteinVolume software ^44^.

|  |  | LoTP | HiT | HiP |
| --- | --- | --- | --- | --- |
| Unit Cell | a (Å) | 39.67 | 39.98 **(+0.8%)** | 39.15 **(-1.3%)** |
|  | b (Å) | 63.51 | 64.49 **(+1.5%)** | 63.45 **(-0.1%)** |
|  | c (Å) | 135.16 | 137.21 **(+1.5%)** | 134.22 **(-0.7%)** |
|  | Cell Volume (Å<sup>3</sup>) | 340527.7 | 353769.9 **(+3.9%)** | 333411.5 **(-2.1%)** |
| Protein | Protein Volume (Å<sup>3</sup>) | 37616.6 | 38054.3 **(+1.2%)** | 37798.3 **(+0.5%)** |

### Alterations to ordered solvent

In addition to these global changes to the crystal lattice, high temperature and pressure have striking effects on the solvation layer surrounding the protein. Compared to LoTP, both HiT and HiP have significantly fewer ordered water molecules (**Fig. 2**). Turning to specific positions, 13 (14.2%) of the HiT waters and 22 (19.3%) of the HiP waters were distinct from any LoTP water (> 2 Å, accounting for crystal symmetry). Of these 35 new positions, only 4 (11.4%) were common to both HiT and HiP.

**Figure 2:**
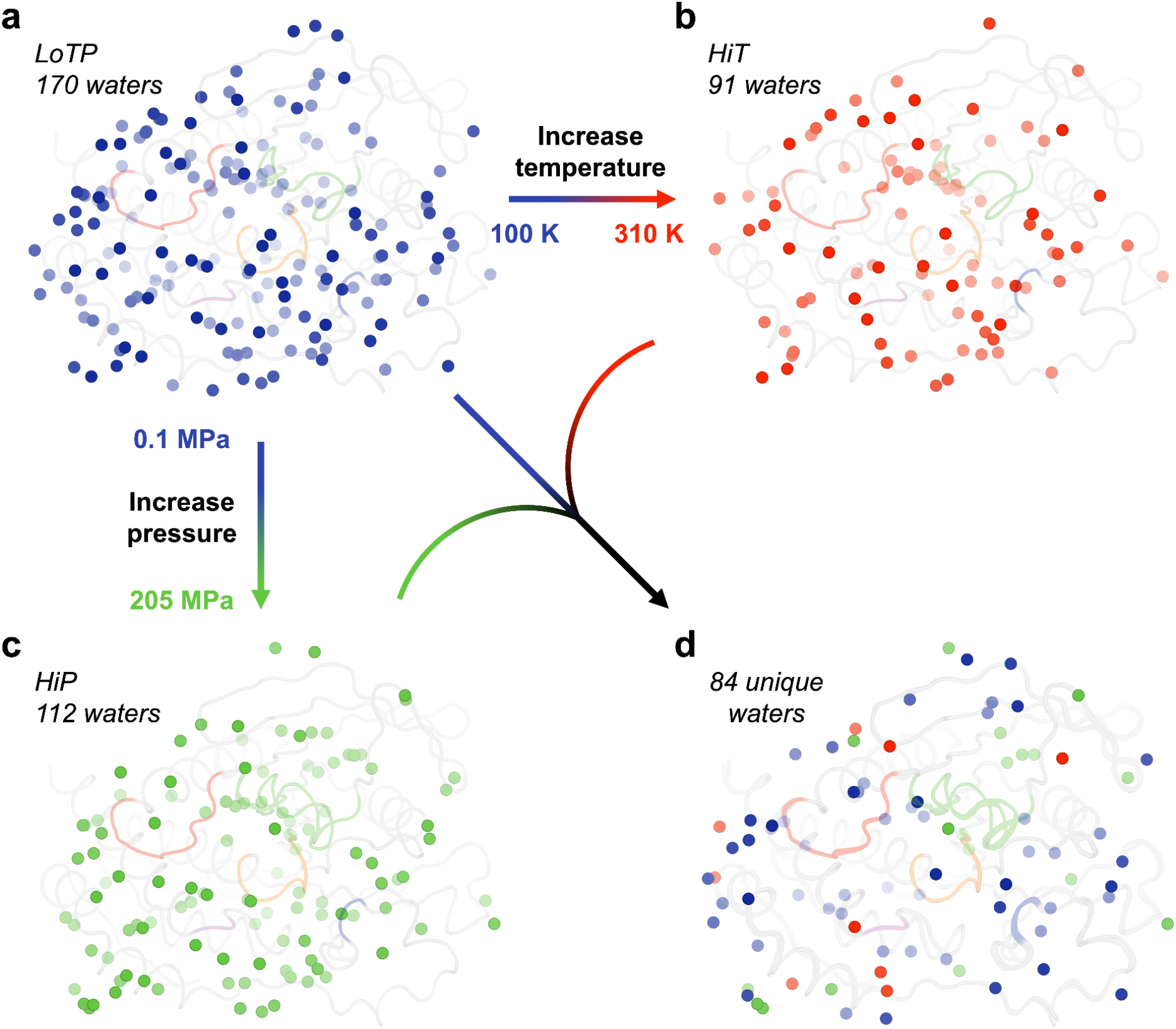
Ordered water molecules are sensitive to temperature and pressure. (a-c) All ordered water molecules at (a) LoTP, (b) HiT, and (c) HiP are shown. (d) Only the waters unique to each structure, i.e. > 2Å from any water in the other two structures. Coloring for active-site loops as in **Fig. 1**.

This suggests that high temperature and pressure do not merely retain a subset of ordered waters, but rather stabilize new water positions, resulting in a distinct pattern of solvation. As shown below, some of these unique waters are located at functional sites in STEP (**Fig. 3**).

**Figure 3:**
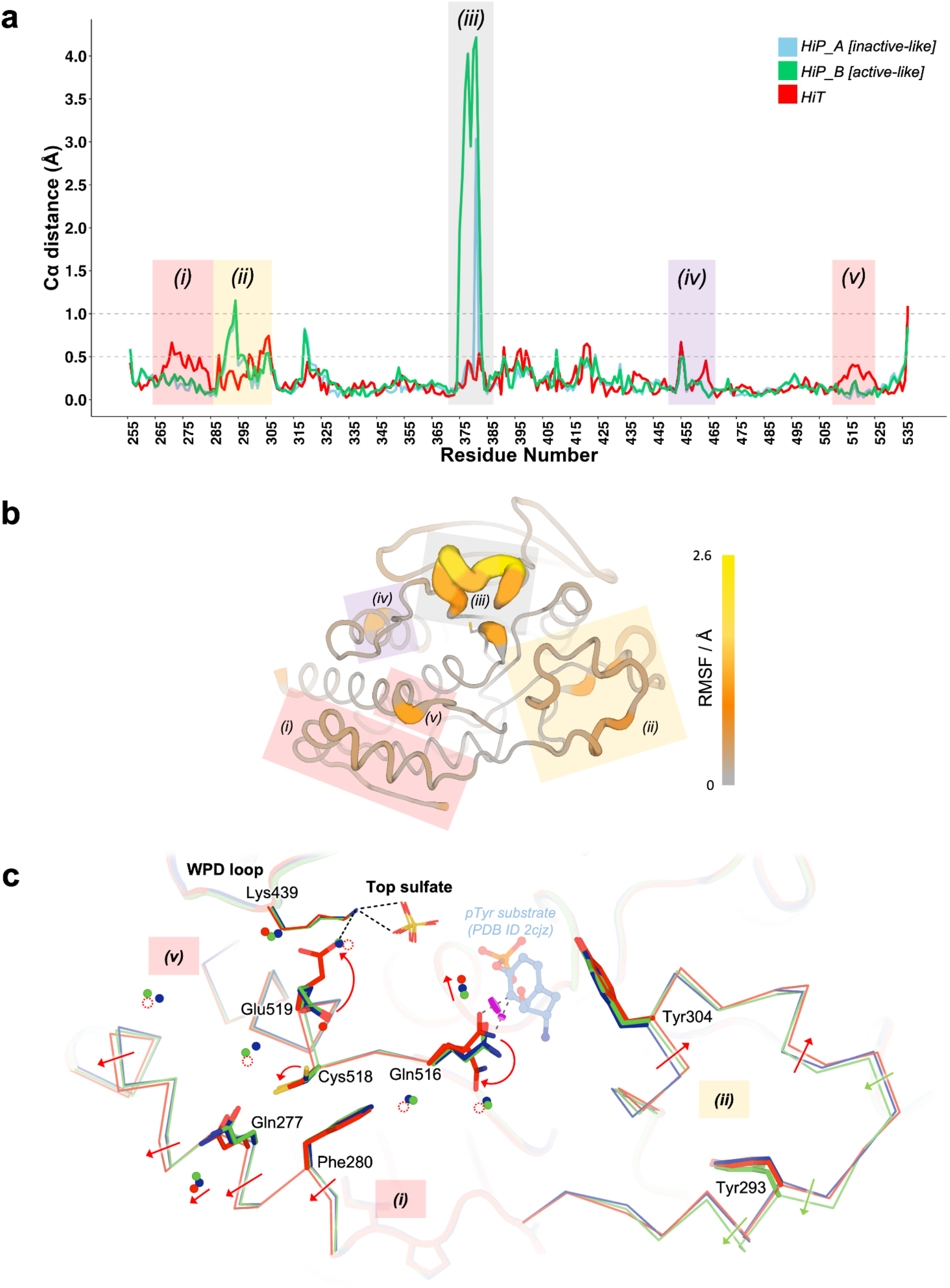
Global backbone displacements due to high temperature vs. pressure. a) Cɑ distances for the HiT and HiP structures relative to the reference LoTP structure are plotted vs. amino acid sequence. The two alternate conformations for the E loop in the HiP structure are separated, although both have high Cɑ distance to LoTP. See also **Supp. Figure 3**. Regions with interesting backbone differences are highlighted; those highlighted with the same color are adjacent in the tertiary structure. b) Structure of STEP with color and cartoon width corresponding to Cɑ root-mean-square fluctuation (RMSF) between our HiT, HiP, and LoTP structures. Same highlighted regions as in (a). c) Zoom-in of active-site area including region (i) (residues 267-282, ɑ1’-ɑ2’ helices), region (v) (residues 515-531, Q loop), and region (ii) (residues 287-306, pTyr loop). Catalytic WPD loop and “top” sulfate are shown nearby. Magenta disks show putative steric clashes between Gln516 and an aligned pTyr substrate from PDB ID 2cjz (not in our structures) which is included for context. See **Fig. 3** for zoom-in of region (iii), and **Supp Fig. 4** for zoom-in of region (iv).

### Global effects on protein conformation

To explore differential effects of high temperature vs. pressure on the protein molecule itself, we examined Cɑ displacements in the HiT and HiP structures relative to the reference LoTP structure. A global plot of this Cɑ distance vs. amino acid sequence (**Fig. 3a**) reveals that most regions are similar in the three structures, with Cɑ distances < 0.3 Å, but several local regions shift relative to the reference structure. These shifts tend to occur either only at high temperature or only at high pressure, suggesting that the protein responds to these different perturbations in distinct ways.

Our structures were obtained in the same crystal form as PDB ID 2bv5, which, like our LoTP structure, is a cryogenic-temperature, ambient-pressure dataset. Cɑ distance analysis shows that for many key regions, 2bv5 is similar to our LoTP structure, whereas our HiT and HiP structures are more different (**Supp. Fig. 3**). Thus the effects of temperature and pressure are generally greater than the variability inherent to determining structures of the same protein in similar conditions by different scientists at different times.

Beyond 2bv5, all other previous human STEP structures were in a different crystal form (same space group but longer a and shorter c axes). These exhibit similar or greater Cɑ distances than do our HiT and HiP structures at several sites in STEP (**Supp. Fig. 3b**). All previous STEP structures were determined at cryogenic temperature and ambient pressure. This indicates that, at least at a gross level, differences in crystal contacts may elicit protein structural variability ^45^ that encompass much of the variability elicited by experimental perturbations such as temperature and pressure. Nonetheless, as shown below, temperature and pressure each induce novel, unique conformational states of STEP.

### Local effects on key structural regions

To explore the basis of these global structural differences, we examined several local areas with distinct conformations in the HiT vs. HiP structures. One local region that responds strongly to pressure -- but not to temperature -- is the E loop (**Fig. 3a** region (iii)). The E loop of PTPs, containing several Glu (E) residues, is located adjacent to the catalytic WPD loop and P loop (**Fig. 1**). Among previous structures of STEP, the E loop exhibited substantial variability (**Supp. Fig. 1, Supp. Fig. 3**). The two main states previously modeled for this loop were the “inactive-like” state in 2bv5 (with an acetylated catalytic Cys472), and the “active-like” state in 6h8r (bound to a distal allosteric small-molecule activator). To validate these previous models, we inspected the electron density maps for all previous STEP crystal structures (7 human, 1 mouse). We determined that all of these structures besides 6h8r were either already modeled with a 2bv5-like conformation, or were unmodeled but could be better explained by a 2bv5-like conformation than by a 6h8r-like conformation. Thus, the “active-like” state of the E loop was only legitimately observed in the allosterically activated structure 6h8r, even though the density was somewhat noisy (**Supp. Fig. 4**).

In contrast to previous STEP structures, our HiP electron density for the E loop, albeit also noisy, is consistent with the presence of both an inactive-like state as in 2bv5 and an active-like state as in 6h8r. We therefore modeled both states as alternate conformations (**Fig. 4a-b**). Deletion of either of these conformations and calculation of omit maps results in positive Fo-Fc difference density peaks for the omitted model (**Fig. 4c-d**), suggesting both are present. The 6h8r-like conformation exists in our HiP structure despite having a different crystal form than 6h8r. By contrast to HiP, our crystallographically isomorphous LoTP and HiT structures are essentially identical to 2bv5 for the E loop. Therefore, high pressure appears to uniquely stabilize a conformation of a key active-site loop that is correlated with an allosterically activated state of human STEP.

**Figure 4:**
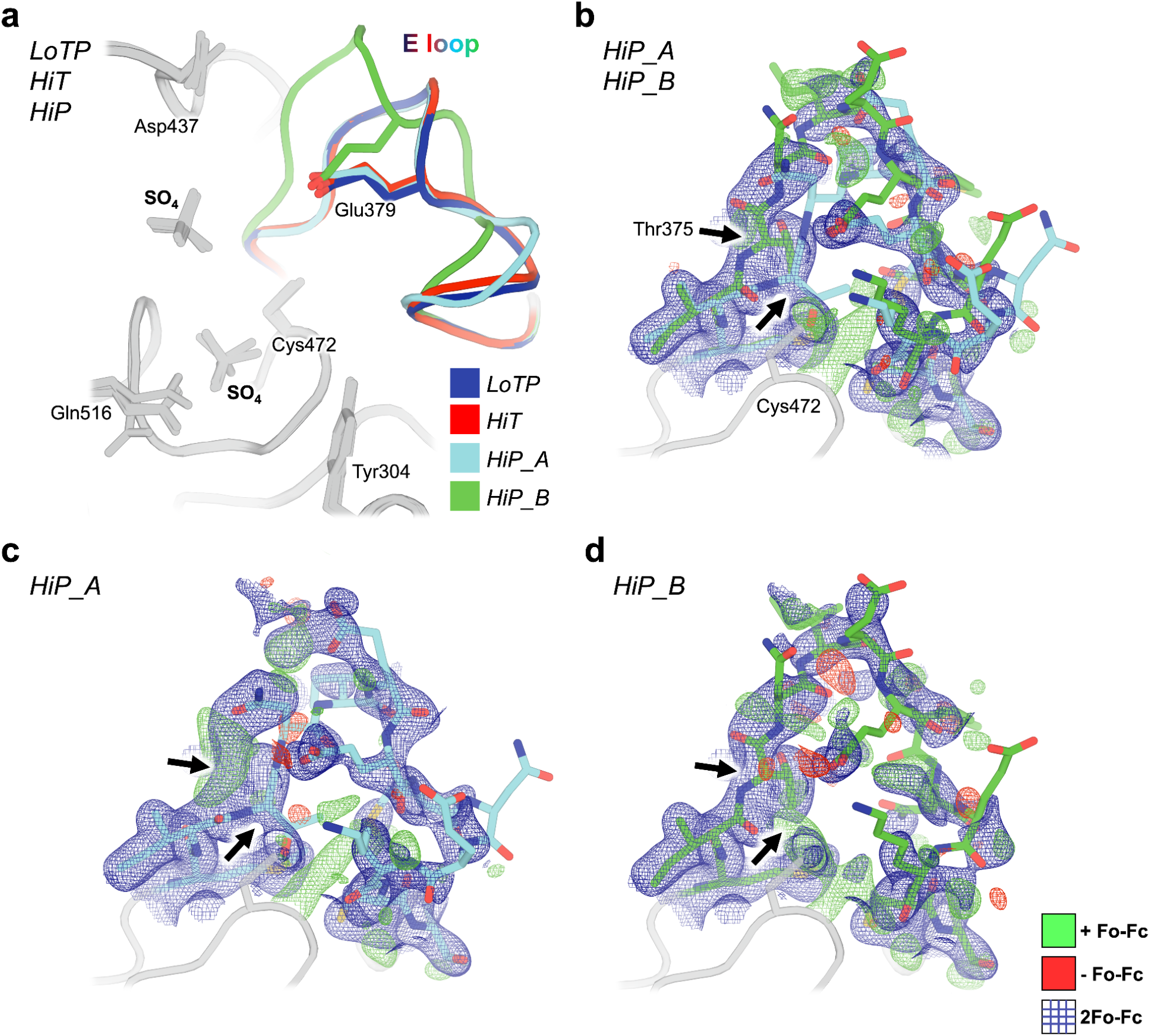
The E loop is reorganized uniquely at high pressure. a) Overlay of all three new STEP structures. The conformation of the E loop is nearly identical at LoT (blue) and HiT (red), but deviates into two distinct conformations at HiP (cyan, green). Glu379 remains within the same conformational space regardless of E-loop conformation. b) HiP dual E loop (cyan, green sticks) with 2Fo-Fc (blue mesh, 1 σ) and Fo-Fc difference (green mesh, 3 σ; red mesh, -3 σ) electron density maps. Thr375 is the first full amino acid where the E loop completely separates into two distinct conformations (black arrows). c) HiP_A conformation of E loop with 2Fo-Fc and Fo-Fc difference electron density maps, omitting the HiP_B state. Arrows indicate Thr375 deviation. d) HiP_B conformation of E loop with 2Fo-Fc and Fo-Fc difference electron density maps, omitting the HiP_A state. Arrows indicate Thr375 deviation.

Another region that responds to pressure is residues 287-306, encompassing the pTyr loop, also known as the substrate-binding loop (SBL) (**Fig. 3a-b** region (ii)). Backbone shifts in this region play a crucial role in defining the depth of the catalytic pocket ^46^. In our models, the backbone of this region, particularly the N-terminal portions, shifts up to ∼1 Å from LoTP to HiP (**Fig. 3c**). In addition, the backbone of the C-terminal portions of this region, corresponding to the pTyr loop itself, shifts by up to 0.75 Å from LoTP to HiT (**Fig. 3c**). Notably, the backbone for the pTyr loop residue Tyr304, whose side chain directly interacts with and helps position the pTyr substrate during catalysis, shifts at both HiP and HiT, yet its side chain remains in place. Overall, these observations suggest a degree of plasticity in the substrate-binding region, which we speculate may help accommodate different pTyr-containing substrates.

In contrast to these regions that respond to pressure, other regions of STEP respond only to temperature. The backbone of the ɑ1’-ɑ2’ helical region near the N-terminus (residues 266-284) shifts by up to ∼0.6 Å at HiT but not HiP (**Fig. 3a-b** region (i)). In addition, the junction between the active-site Q loop and the ɑ6 helix (residues 514-524) shifts by up to ∼0.4 Å, also at HiT but not HiP (**Fig. 3a-b** region (v)). These new HiT conformations differ not only from our HiP and LoTP structures, but also from the only previous STEP structure with the same crystal form, 2bv5, which was at cryogenic temperature (**Supp. Fig. 3a**).

These backbone shifts are coupled to other notable changes to solvation and side-chain conformational ensembles (**Fig. 3c**). First, several ordered waters are liberated from the interface between ɑ1’-ɑ2’ and the Q loop only at HiT, illustrating a complex interplay between protein and solvent structure. Also, in concert with the Q loop backbone shift in this interface, the side chain of Cys518 (from the Q loop) switches from two rotamers to one. The disappearance of the alternate rotamer for Cys518 eliminates a hydrogen bond to the adjacent Glu519, causing the latter to switch to a new rotamer (see also **Fig. 5b**). The new Glu519 rotamer engages in a previously unseen interaction with Lys439 from the catalytic WPD loop, which coordinates the top sulfate (**Fig. 3c**). This novel conformation of Glu519 is not present in any previous STEP structures: it is unique to our HiT structure.

**Figure 5:**
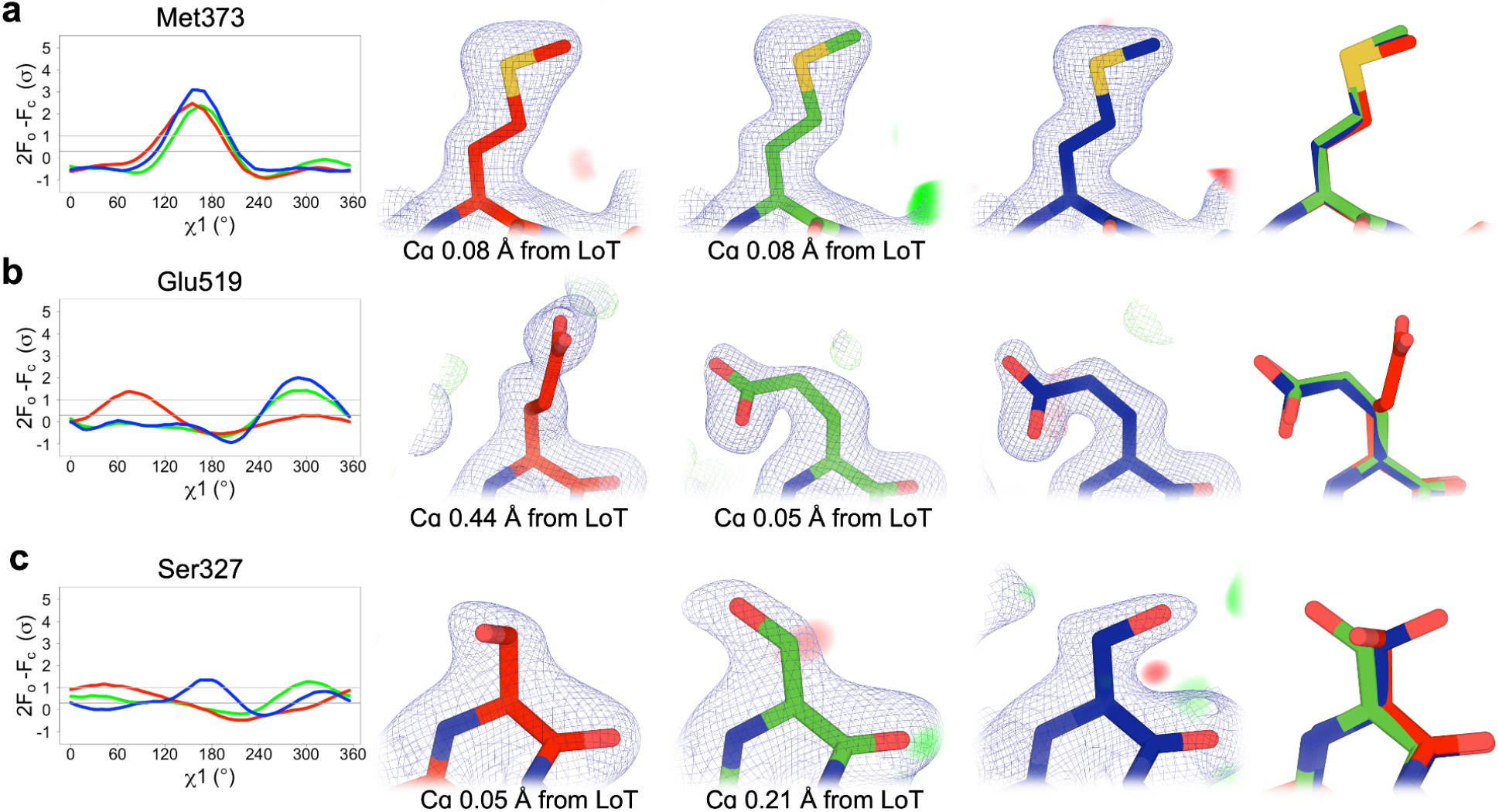
Examples of different side-chain conformations at high temperature and/or pressure. For each example residue, the following panels are shown: (*Left*) Overlaid Ringer curves for our three datasets ^4^. (*Middle*) Our three structures with 2Fo-Fc (contoured at 1 σ) and Fo-Fc (contoured at ±3 σ) density maps. *(Right)* Our three structures overlaid. LoTP in blue, HiT in red, HiP in green. Examples: a) Met373 has the same χ1 peak (*t*, near 180°) for LoTP, HiT, and HiP. b) Glu519 has similar χ1 peaks for LoTP and HiP (*m*, near -60°), but a different peak for HiT (*p*, near +60°). c) Ser327 has different χ1 peaks for LoTP (*t*), HiT (*p*), and HiP (*m*). Χ^1^ rotamer nomenclature from ^48^.

The Q loop backbone shift is also correlated with an alternate side-chain rotamer for Gln516 (**Fig. 3c**) that has only been seen in two previous structures: bound to a pTyr substrate (2cjz) (**Supp. Fig. 2c**), and bound to a distal allosteric activator (6h8r) (**Supp. Fig. 5**). In particular, this new rotamer avoids what would otherwise be a steric clash with the pTyr substrate, which binds immediately adjacent to Gln516 (**Fig. 3c**). These observations suggest that HiT may capture an active-like conformation of STEP, even in the apo form, that is more compatible with formation of the Michaelis complex.

Interestingly, Gln516 is immediately adjacent to Ile515, which is the only residue in STEP to have an unavoidable but real Ramachandran outlier -- consistent with previous observations that validated, geometrically strained residues, while rare, occur preferentially at active sites ^47^.

In the context of the crystal lattice, ɑ1’-ɑ2’ also abuts the distal S loop (residues 462-465), parts of which shift by > 0.4 Å at HiT but not HiP (**Supp. Fig. 5**). Interestingly, the S loop forms part of the binding site for a class of allosteric small-molecule activators that are unique to STEP ^39^ (**Supp. Fig. 5**). This coincidence of temperature-sensitive regions in 3D space suggests that subtle lattice expansion at elevated temperature can allow a protein “breathing room” to adopt subtly different conformations, including at functionally important regions.

### Widespread changes to torsion angles

Complementing this analysis of specific regions, we examined in detail how temperature vs. pressure affected conformations throughout the entirety of the STEP catalytic domain, using torsion angles in several ways. First, we performed Ringer analysis for each side chain in each structure by rotating around the Cα-Cβ vector (χ1 torsion angle) and measuring the 2Fo-Fc electron density value at each possible γ heavy atom position ^4^. For each residue, we then calculated a correlation coefficient (CC) between Ringer curves for each pair of datasets ^15^. Relative to the “reference” LoTP dataset, a substantial number of residues had low CC for either HiT or for HiP (**Supp. Fig. 6**), suggesting differences in side-chain conformational ensembles due to these perturbations. For example, 18 (6.4%) residues had CC < 0.5 in HiT, and 16 (5.7%) residues had CC < 0.5 in HiP. Excluding the flexible E loop (residues 375-383), 18 (6.6%) residues had CC < 0.5 in HiT, and 13 (4.7%) residues had CC < 0.5 in HiP. If high temperature vs. high pressure had similar structural effects, a similar set of residues would be expected to have low CC for both HiT and HiP (each relative to LoTP). However, relatively few residues fall into this category (purple bars in **Supp. Fig. 6**), suggesting that temperature vs. pressure are complementary perturbations that affect different areas of the protein.

To validate these quantitative Ringer results, we examined the models and density maps in detail for several examples. For most residues, the Ringer curves are indeed similar across all three datasets (**Fig. 5a**). For other residues, by contrast, the curves differ in one or more datasets, indicating perturbation-induced changes to side-chain conformations. For example, Glu519 adopts the same χ1 rotamer for LoTP and HiP, but a different χ1 rotamer at HiT (**Fig. 5b**; see also **Fig. 3c**), involving a ∼0.4 Å backbone shift (**Fig. 3a**). By contrast, Ser327 adopts different primary χ1 rotamers for LoTP, HiT, and HiP (**Fig. 5c**). Some residues had distinct Ringer curves at HiP relative to LoTP and HiT (**Supp. Fig. 7**) but were associated with distinct backbone positions of the E loop that occurred only at HiP (**Fig. 3a, Fig. 4**).

As the Ringer curves above only account for the first side-chain torsion angle (χ1), we also compared rotamer names, which account for all side-chain torsion angles ^48^ (see Methods). Excluding the flexible E loop, relatively few residues had different rotamers as alternate conformations in the same model: 6 for LoTP, 5 for HiT, and 5 for HiP. However, compared to LoTP, 43 residues (18%) had a different rotamer in HiT, and 33 residues (14%) had a different rotamer in HiP. Thus, by contrast to only the side-chain “base” as measured by Ringer, temperature and pressure both have greater effects on the overall conformations of side chains, stabilizing distinct energy basins. Moreover, of the residues that differed from LoTP, 30 were unique to either only HiT or only HiP, indicating distinct conformational effects from temperature vs. pressure.

Finally, beyond just torsion angles for individual side chains, we explored whether many torsion angles distributed throughout the protein structure may undergo correlated changes in response to temperature vs. pressure. A recent tool called RoPE showed that linear combinations of backbone and side-chain torsion angles in a reduced-dimensionality space can help reveal new insights into the key differences between sets of structural models ^40^. Using RoPE analysis, we examined our three new structures relative to all previous STEP structures (**Fig. 6**), leading to several interesting observations.

**Figure 6:**
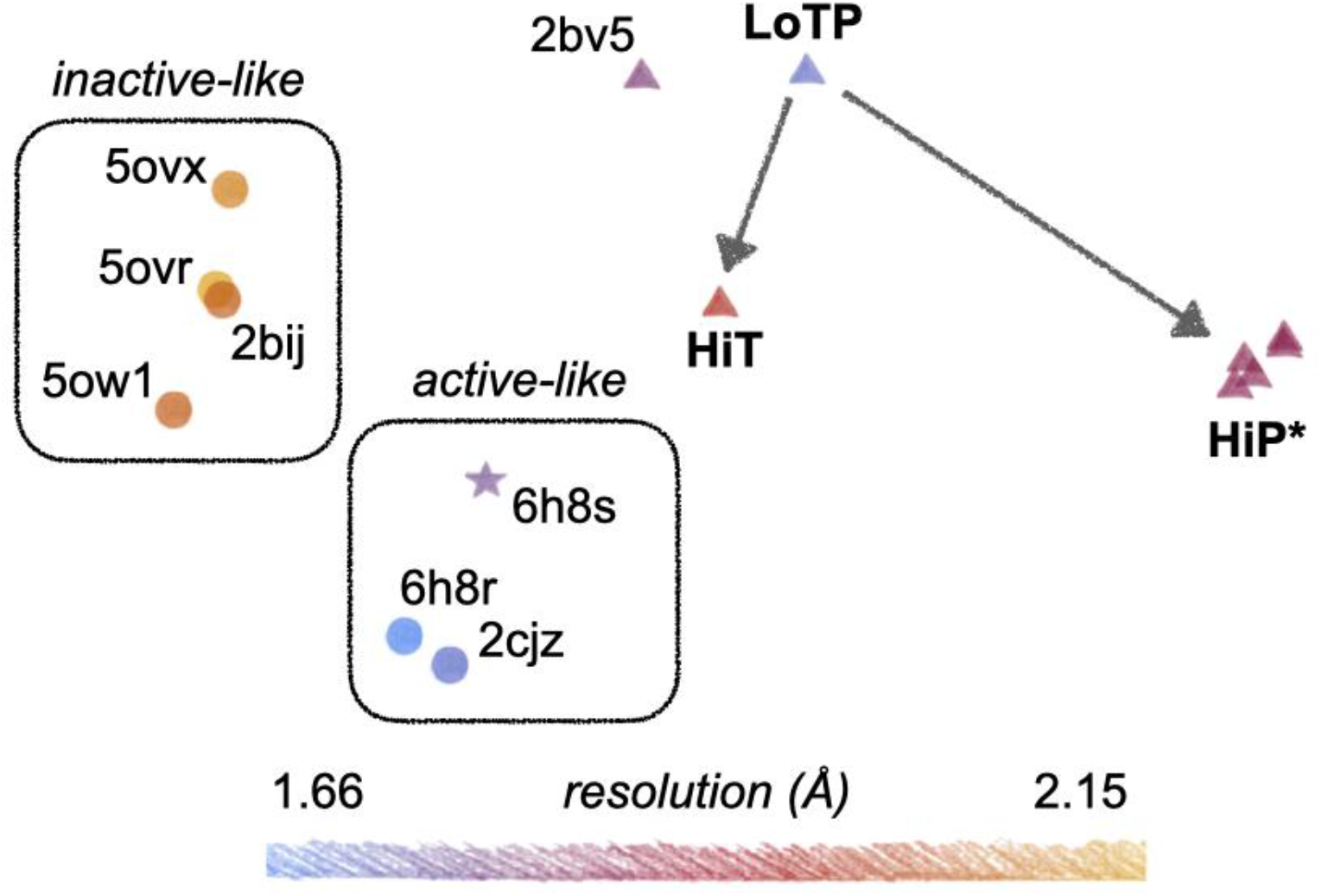
Dimensionality reduction in torsion-angle space reveals clustering based on several factors including temperature vs. pressure. Our new structures (LoTP, HiT, HiP) are shown relative to all previous STEP structures from the PDB in RoPE reduced-dimensionality torsion-angle space ^40^. Horizontal and vertical axes correspond to different combinations of the top principal component analysis (PCA) modes. Apparent inactive-like vs. active-like clusters are highlighted. Different icon shapes indicate distinct crystal forms (unit cell parameters). Resolution is shown by color. Arrows indicate the effects of high temperature vs. pressure relative to our reference structure. HiP* indicates several HiP models with the E loop prepared in different ways; see Methods. 6h8s: mouse STEP; all other structures: human STEP. All models were prepared for RoPE with PDB-REDO ^49^ to ensure consistent treatment ^40^.

First, the STEP structures generally cluster based on resolution, as noted previously for other proteins ^40^, with our new structures at intermediate-to-high resolution compared to prior structures. Second, each set of structures with a consistent crystal form clusters together: (i) our new structures plus 2bv5, (ii) most remaining structures, and (iii) the mouse STEP structure 6h8s. Third, most of the previous structures segregate into an active-like cluster (either allosterically activated or bound to a substrate peptide) or an inactive-like cluster (bound to orthosteric inhibitors), indicating that subtle signatures of the protein’s functional state are embedded in torsion-angle space. The allosterically activated structure 6h8s is nearest to other active-like structures, despite it being mouse-derived (91% sequence identity to human STEP) and having a unique crystal form; hence, signatures of the protein’s inherent functional state appear to persist in this space despite differences in amino acid sequence and crystal lattice.

Fourth, whereas our LoTP structure is near the most analogous previous structure (2bv5) in torsion-angle space as expected, our HiT and HiP structures move in distinct directions from this reference point (**Fig. 6**). Notably, HiT moves toward the active-like structures, whereas HiP moves away from all known STEP structures. This is despite only HiP featuring a conformation of the E loop resembling the allosterically activated structure 6h8r (**Fig. 4, Supp. Fig. 4**), but consistent with only HiT featuring side-chain and backbone conformations of the active-site Q loop in a putatively active-like state (**Fig. 3c, Fig. 5b**). HiP models with significantly different E-loop conformations and prepared for analysis in different ways have similar positions in torsion-angle space, confirming that RoPE analysis highlights structurally distributed as opposed to localized features. Relative to LoTP, the coordinated torsion-angle changes in HiT and HiP detected by RoPE visually correspond to “hinging” of the first half of the primary structure (initial α-helices + loops) relative to the second half (β-sheet + α-helical bundle), albeit with apparently meaningful differences between them given their large separation in RoPE space. Overall, these results indicate that although high pressure induces an active-like conformation locally in the E loop, high temperature induces a more global, distributed active-like state of the protein.

## Discussion

While temperature is growing in use as an experimental perturbation in macromolecular crystallography, pressure has received less attention for such applications. Here we show that both temperature and pressure enact distinct and significant effects on the conformational ensemble of STEP, not only globally but also locally at several key functional areas.

In our structures, high temperature increases both unit cell volume and protein molecular volume (**Table 2**) as seen previously ^9^. High pressure decreases unit cell volume as seen previously ^27,50,51^, yet still slightly increases protein molecular volume (**Table 2**). Thus, intriguingly, when subjected to pressure, the STEP protein molecule itself slightly expands, even as its environment is compressed. This differs from previous high-pressure crystal structures of other proteins with a decreased protein molecular volume ^27,29,31^, but agrees with a high-pressure NMR solution structure with a slightly increased protein molecular volume, corresponding to negative compressibility ^52^. Our surprising, counterintuitive result suggests that the mechanisms by which pressure impacts the conformational landscapes of different proteins are complex and potentially context-sensitive.

Although many of the structural changes we see could be considered small, it is important to remember that sub-angstrom shifts can be directly relevant to protein function ^26^. This is consistent with our RoPE results, in which HiT vs. HiP have very distinct characteristics despite the overall structures being apparently similar. Crucially, the difference in the E loop at HiP does not dominate the signal (see **Fig. 6** and Methods), indicating that the differences between high temperature vs. pressure are driven by smaller, subtler conformational changes distributed throughout the tertiary structure.

The largest conformational changes we observe in STEP are in the E loop (**Fig. 4a**), a conserved loop in PTPs that plays a critical role in regulation ^53^. Only at HiP do we see evidence in the electron density for a dual-conformation E loop (**Fig. 4b-d**). Both conformations were individually evident in previous structures of STEP with different chemical modifications or allosteric ligands (**Supp. Fig. 4**). Our data indicate that applying a physical perturbation (pressure) is sufficient to induce these conformations to coexist in a single crystal of the apo protein, which has implications for accessing excited states of other proteins.

Beyond the E loop, we observe new conformations not captured in previous structures of STEP. For instance, only at HiT, we see Glu519 of the active-site Q loop adopt a new side-chain rotamer that engages in a novel interaction with Lys439. Notably, Lys439 follows the WPD sequence, forming a WPDQK sequence. Recently, a conserved PDFG motif was proposed to underlie the ability of the WPD loop to toggle between discrete open vs. closed states in PTPs ^54^. However, the corresponding residues in STEP are PDQK -- and, as revealed by our HiT structure, the final K (Lys439) engages with the Q loop nearby. Perhaps not coincidentally given these idiosyncratic features, the WPD loop of STEP has not been observed in the usual open or closed states as with most other PTPs, but only in the “atypically open” state ^38,41^. Together, these observations suggest that the STEP active site does not adhere to expectations from the rest of the PTP family, and points to specific amino acids and conformations that may encode its unusual behavior. We speculate that these unique structural properties of STEP likely underlie its substantially lower catalytic activity relative to other PTPs like PTP1B ^11,39,55^ and may be related to its unique physiological roles in neuronal development ^56^.

It is plausible that this unresponsive, atypically open state could be modulated by binding of regulators or alterations in the cellular environment, creating a way to regulate catalysis. In this light, we observe two sulfates simultaneously bound within the active site. The biological significance of this observation for STEP is not immediately clear. It is likely that binding of sulfate-like moieties in the top site, as in several orthosteric inhibitors (**Supp. Fig. 2b**), blocks closure of the WPD loop, effectively wedging it atypically open (**Supp. Fig. 2e**). However, even with the top site free and substrate bound only to the bottom site, the WPD loop still remains atypically open in crystals ^38^ (**Supp. Fig. 2c**).

Removing all tightly bound molecules from the active site of STEP in future crystallographic studies could provide more definitive answers about the conformational landscape of this functionally critical but unusual catalytic loop.

Previously, based on computational simulations, a small-molecule allosteric activator for STEP was reported to enact its effects via a pair of allosteric pipelines ^39^. We do not observe obvious shifts along these pathways at high temperature or pressure. However, we do observe shifts in the activator binding pocket itself (**Supp. Fig. 5**). Although subtle, these conformational shifts may be sufficient to influence ligand-binding energetics, and therefore may aid structure-based drug design efforts to improve upon the relatively weak reported activator.

Beyond the reported allosteric activator site, we also observe perturbation-sensitive shifts at other known or putative ligand binding sites. First, at LoTP and HiP, an ordered glycerol molecule is bound near α2’ and the Q loop. By contrast, at HiT, ordered waters are present instead, and α1’-α2’ and the Q loop undergo conformational shifts (**Fig. 3c**). Importantly, all three structures are from crystals treated with similar glycerol-containing cryoprotectant solutions. It is thus plausible that crystal cryocooling induces the glycerol to bind ^14^, preventing nearby conformations with potential functional relevance seen at physiological temperature (**Fig. 3c, Fig. 6**). Second, in the paralogous PTP SHP2, the α1’-α2’ region helps form the binding site for the potent allosteric inhibitor SHP099, although the mechanism also involves additional domains ^57^. The corresponding α1’-α2’ region in STEP is not known to be allosteric. However, the subtle but coordinated conformational changes we observe here at physiological temperature raise the enticing possibility that some aspects of the allosteric capacity demonstrated in SHP2 are also present in STEP, and perhaps even other PTPs.

In general, allostery in STEP remains poorly understood, hindering efforts to elucidate this important protein’s endogenous regulatory mechanisms and to develop specific allosteric modulators. To address this important gap, several approaches should be considered. First, exploiting different crystal forms, including those in the allosteric-activator-bound structures for human and mouse STEP ^39^, may provide new windows into conformational mobility otherwise masked by crystal contacts. Second, higher pressures than those reported here ^32^ may enable access to additional excited states. Third, X-ray diffraction at high pressure and physiological temperature simultaneously has the potential to reveal unique aspects of conformational landscapes not evident from a single perturbation alone. More broadly, the avant-garde crystallographic and computational methods outlined here should prove useful tools to investigate allosteric mechanisms in a variety of other proteins, including but not limited to other PTP family members that also exhibit an atypically open WPD loop such as LYP ^58^.

Overall, the work reported here is consistent with the notion that proteins sample conformations from a multifaceted energy landscape, and that different physical perturbations such as temperature and pressure can access distinct, complementary features of this landscape, thus opening doors to elucidating fundamental connections between protein structural dynamics and function.

## Methods

### Molecular biology

A plasmid containing the catalytic domain [258–539] of STEP (PTPN5) with an N-terminal 6xHis & TEV cleavage site was obtained via Addgene from Nicola Burgess-Brown (Addgene plasmid #39166; http://n2t.net/addgene:39166; RRID:Addgene_39166). This was transformed into BL21(DE3) Rosetta2 (pRARE2) cells (MilliporeSigma). The sequence of the insert was independently verified using Sanger sequencing, with standard T7 promoter primers.

### Protein expression

In all steps, the antibiotics chloramphenicol (Cam) and ampicillin (Amp) were used to maintain selection at working concentrations of 30 μg/mL and 100 μg/mL respectively. Previously transformed cells from glycerol stocks were plated on an LB-Agar + Amp + Cam plate and incubated overnight at 37°C. Individual colonies were picked and grown up overnight at 18°C in LB + Amp + Cam starter cultures (10 mL), shaking at 180 rpm. This starter culture was then added into baffled flasks containing 1 L of LB+Amp+Cam media, and incubated to OD 0.6–0.8 at 37°C, with shaking at 180 rpm. Expression was then induced by adding IPTG to a final concentration of 0.2 mM; cultures were then incubated overnight at 18°C, shaking at 180 rpm, before cells were harvested by centrifugation at 3000 rpm for 45 minutes, snap frozen in liquid N_2_ and stored at -80°C.

### Protein purification

Frozen cellets (cell pellets) were thawed on ice, then 30 mL lysis buffer (50 mM HEPES pH 7.5, 500 mM NaCl, 5 mM imidazole, 5% v/v glycerol, 2 mM DTT) was added. One Pierce EDTA-free protease inhibitor mini-tablet per cellet was also added, and resuspended in a vortexer. Cells in the slurry were then lysed by 3 passages through a cell homogenizer (Avestin) operating with 1000 bar peak. Lysate was then centrifuged for 45 min at 50000 g to spin down the cell fragments. The supernatant was filtered through a 0.22 μm filter to remove final cell debris.

A 5 mL Ni-NTA column (Cytiva) was equilibrated in freshly prepared low-imidazole buffer (50 mM HEPES pH 7.5, 500 mM NaCl, 30 mM imidazole, 5% v/v glycerol, 2 mM DTT). The lysate supernatant was applied to this column, washed with 2 column volumes (CV) of low-imidazole buffer, then gradient-eluted over 10 CV to 100% high-imidazole buffer (50 mM HEPES pH 7.5, 500 mM NaCl, 500 mM imidazole, 5% v/v glycerol, 2 mM DTT), collecting in 5 mL fractions. The STEP-containing fractions eluted around the 40% gradient mark were collected, concentrated using a 15 mL Centriprep 10K spin-concentrator (Millipore) to a final volume of 5 mL, and filtered through a syringe-mount 0.22 μm filter to remove unidentified precipitate.

A Sephadex 20/200 column (Cytiva) was equilibrated with 2 CV of SEC buffer (50 mM HEPES pH 7.5, 500 mM NaCl, 5 % v/v glycerol, 2 mM DTT). The concentrated, filtered Ni-binding fraction was injected onto a 5 mL loop, loaded onto the column, and fractionated over 2 CV, collecting 1 mL fractions. Two peaks were observed, and the fractions corresponding to the largest, STEP-containing peak were pooled.

A HiTrap Q HP anion-exchange column (Cytiva) was equilibrated with 2 CV of low-salt buffer (50 mM HEPES pH 7.5, 10 mM DTT). The pooled peak from size-exclusion chromatography was diluted to a final volume of 100 mL by addition of low-salt buffer, and filtered through a 0.22 μm bottle-top vacuum filter (Celltreat). This was then applied to the Q column, washed with 2 CV of low-salt buffer, and then gradient-eluted over 5 CV to 100% high-salt buffer (50 mM HEPES pH 7.5, 1000 mM NaCl, 10 mM DTT) collecting 5 mL fractions. A single STEP-containing peak was collected at 40% gradient mark.

This final STEP protein was concentrated in Centriprep 10K spin-concentrators to 3 mL volume, and then further concentrated in Amicon 10K spin-concentrators to a final concentration of 10 mg/mL, as measured by Nanodrop, and used fresh as the protein sample in crystallography. The identity of STEP vs. other proteins/contaminants was confirmed using SDS-PAGE gels at each step of the purification.

### Crystallization and crystal preparation

Precipitant well solution (30% PEG 3350, 200 mM Li_2_SO_4_, 100 mM bis-tris pH 5.65) was prepared fresh. A Mosquito (SPT Labtech) was used to prepare 96-well 3-drop Intelliplate Low-profile (Art Robbins Instruments) plates. 80 μL well solution was placed into the reservoir. Three 1 μL drops at a protein concentration of 10 mg/mL were placed per well, using 2:1, 1:1, and 1:2 ratios of well solution to protein sample. Crystallization drops were incubated at room temperature. Crystals nucleated within 3 days, mostly in 1:1 droplets, and grew over a week to around 80 × 80 × 20 μm.

For the ambient-pressure low-temperature 100 K (LoTP) dataset, the crystal was soaked in cryoprotectant (mother liquor + 15% v/v glycerol), and cryocooled with liquid nitrogen.

For the high-pressure (205 MPa) cryogenic-temperature dataset (HiP), the crystal was looped in a 100 μm loop, soaked in cryoprotectant (mother liquor + 15% v/v glycerol), and placed in a capillary with cryoprotectant at the end of the tube for shipping to CHESS. At CHESS, the capillary was removed, and the crystal was coated in NVH oil. The crystal was pressurized for 20 min at 205 MPa, cryocooled with liquid nitrogen under pressure, and stored under liquid nitrogen thereafter.

High-temperature diffraction required larger crystals, and so were prepared in Nextal EasyXtal 15-well hanging-drop trays. Precipitant well solution was prepared with the same composition as above (pH 5.5). 400 μL well solution was placed into the reservoirs. Three 3 μL drops at a protein concentration of 10 mg/mL were placed per well, using 1:1 ratios of well solution to protein sample each.

Crystallization drops were incubated at room temperature. Crystals nucleated within 3 days, and grew over a week to around 140 × 70 × 40 μm.

For the high-temperature ambient-pressure 310 K (HiT) dataset, the crystal was soaked in cryoprotectant (mother liquor + 15% v/v glycerol), coated in NVH oil, and placed in a capillary with cryoprotectant at the end of the tube for shipping to CHESS.

### X-ray data collection

All X-ray diffraction datasets were collected at the ID7B2 (FlexX) beamline for macromolecular X-ray science at the Cornell High Energy Synchrotron Source (MacCHESS), Ithaca, New York, USA, using an X-ray beam energy of 12 keV and corresponding wavelength of 1.033 Å. The LoTP dataset was collected using beam dimensions of 30 × 20 μm, flux of 5x10^11^ ph/s, rotation rate of 2°/s, and no translation. The HiT dataset was collected using beam dimensions of 30 × 20 μm, flux flux of 1.6x10^10^ ph/s, rotation rate of 10°/s, and translation (i.e. helical/vector data collection) along the length of the approximately 140 × 70 × 40 μm crystal. The HiP dataset was collected using beam dimensions of 100 × 100 μm, flux of 2x10^10^ ph/s, rotation rate of 1°/s, and no translation.

### X-ray data reduction and modeling

Data reduction and modeling was performed similarly for all three datasets, with the data reduction pipeline DIALS ^59^. The LoTP dataset was trimmed to the first 130 (out of 180) frames due to increased ice inclusions in later frames. Resolution cutoffs were determined automatically by DIALS based on CC1/2 ^60^. Molecular replacement was performed via Dimple ^61^, with subsequent refinement performed using REFMAC ^62^ and phenix.refine ^63^, with models manually adjusted between rounds of refinement using COOT ^64^. Hydrogens were added using phenix.ready_set ^65^. X-ray/stereochemistry weight, X-ray ADP weight, and occupancies were all refined and optimized during the final rounds of refinement. Model validation statistics were generated using MolProbity ^66^. Data collection and refinement statistics can be found in **Table 1**.

### Model analysis

Cɑ distances between structures were calculated using VMD ^67^. Rotamer names were calculated using phenix.rotalyze ^66^ based on the latest rotamer distributions from MolProbity ^68^.

Protein volumes were calculated using the ProteinVolume software ^44^. Values for the “total volume” output were nearly identical whether waters were included or not, and were similar (conclusions did not change) when state A vs. state B of the HiP structure were analyzed.

Ringer ^4^ was run on models that only contained a single conformation for each residue, with alternate conformations removed using phenix.pdbtools. For HiP, each E-loop conformation was treated individually. The single-conformer models each underwent multiple cycles of refinement using phenix.refine. The refined models and maps were then used as input to Ringer. Plots were generated using *ggplot2* and *ggbreak*.

For RoPE analysis, all structures were pre-processed with PDB-REDO ^49^ to ensure consistent treatment, as recommended ^40^. The different HiP points in **Fig. 6** correspond to different preparations of the model: run PDB-REDO for deposited model; extract each state, run PDB-REDO, then set all occupancies to unity; or extract each state, set all occupancies to unity, then run PDB-REDO.

## Acknowledgments

DAK is supported by NIH R35 GM133769.

We thank Helen Ginn for help with RoPE analysis, members of the CUNY Advanced Science Research Center (ASRC) Structural Biology Initiative (SBI) for helpful discussions, Akshay Raju and Shivani Sharma for help with PTP bioinformatics, and Marian Szebenyi for help with arranging our X-ray beamtime.

This work is based upon research conducted at the Center for High Energy X-ray Sciences (CHEXS), which is supported by the National Science Foundation under award DMR-1829070, and the Macromolecular Diffraction at CHESS (MacCHESS) facility, which is supported by award 1-P30-GM124166-01A1 from the National Institute of General Medical Sciences, National Institutes of Health, and by New York State’s Empire State Development Corporation (NYSTAR).

## References

1. Frauenfelder, H., Sligar, S. G. & Wolynes, P. G. The energy landscapes and motions of proteins. Science 254, 1598–1603 (1991).

2. Henzler-Wildman, K. & Kern, D. Dynamic personalities of proteins. Nature 450, 964–972 (2007).

3. Xie, T., Saleh, T., Rossi, P. & Kalodimos, C. G. Conformational states dynamically populated by a kinase determine its function. Science 370, p(2020).

4. Lang, P. T. et al. Automated electron-density sampling reveals widespread conformational polymorphism in proteins. Protein Sci. 19, 1420–1431 (2010).

5. Wankowicz, S. A., de Oliveira, S. H., Hogan, D. W., van den Bedem, H. & Fraser, J. S. Ligand binding remodels protein side-chain conformational heterogeneity. Elife 11, p(2022).

6. Fraser, J. S. et al. Hidden alternative structures of proline isomerase essential for catalysis. Nature 462, 669–673 (2009).

7. Keedy, D. A. Journey to the center of the protein: allostery from multitemperature multiconformer X-ray crystallography. Acta Crystallogr D Struct Biol 75, 123–137 (2019).

8. Fischer, M. Macromolecular room temperature crystallography. Q. Rev. Biophys. 54, e1 (2021).

9. Fraser, J. S. et al. Accessing protein conformational ensembles using room-temperature X-ray crystallography. Proc. Natl. Acad. Sci. U. S. A. 108, 16247–16252 (2011).

10. Keedy, D. A. et al. Crystal cryocooling distorts conformational heterogeneity in a model Michaelis complex of DHFR. Structure 22, 899–910 (2014).

11. Keedy, D. A. et al. An expanded allosteric network in PTP1B by multitemperature crystallography, fragment screening, and covalent tethering. Elife 7, p(2018).

12. Ebrahim, A. et al. The temperature-dependent conformational ensemble of SARS-CoV-2 main protease (Mpro). IUCrJ 9, p(2022).

13. Fischer, M., Shoichet, B. K. & Fraser, J. S. One Crystal, Two Temperatures: Cryocooling Penalties Alter Ligand Binding to Transient Protein Sites. Chembiochem 16, 1560–1564 (2015).

14. Skaist Mehlman, T. et al. Room-temperature crystallography reveals altered binding of small-molecule fragments to PTP1B. Elife 12, p(2023).

15. Keedy, D. A. et al. Mapping the conformational landscape of a dynamic enzyme by multitemperature and XFEL crystallography. Elife 4, p(2015).

16. Doukov, T., Herschlag, D. & Yabukarski, F. Instrumentation and experimental procedures for robust collection of X-ray diffraction data from protein crystals across physiological temperatures. J. Appl. Crystallogr. 53, 1493–1501 (2020).

17. Ebrahim, A. et al. Resolving polymorphs and radiation-driven effects in microcrystals using fixed-target serial synchrotron crystallography. Acta Crystallogr D Struct Biol 75, 151–159 (2019).

18. Fischer, M., Coleman, R. G., Fraser, J. S. & Shoichet, B. K. Incorporation of protein flexibility and conformational energy penalties in docking screens to improve ligand discovery. Nat. Chem. 6, 575–583 (2014).

19. Bradford, S. Y. C. et al. Temperature artifacts in protein structures bias ligand-binding predictions. Chem. Sci. 12, 11275–11293 (2021).

20. Cavender, C. E. et al. Structure-Based Experimental Datasets for Benchmarking of Protein Simulation Force Fields. arXiv [q-bio.BM] (2023).

21. Kurpiewska, K. & Lewiński, K. High pressure macromolecular crystallography for structural biology: a review. Open Life Sciences 5, 531–542 (2010).

22. Fourme, R., Girard, E. & Akasaka, K. High-pressure macromolecular crystallography and NMR: status, achievements and prospects. Curr. Opin. Struct. Biol. 22, 636–642 (2012).

23. Dhaussy, A.-C. & Girard, E. Functional Sub-states by High-pressure Macromolecular Crystallography. in High Pressure Bioscience: Basic Concepts, Applications and Frontiers (eds. Akasaka, K. & Matsuki, H.) 215–235 (Springer Netherlands, 2015).

24. Roche, J., Royer, C. A. & Roumestand, C. Exploring Protein Conformational Landscapes Using High-Pressure NMR. Methods Enzymol. 614, 293–320 (2019).

25. Xu, X., Gagné, D., Aramini, J. M. & Gardner, K. H. Volume and compressibility differences between protein conformations revealed by high-pressure NMR. Biophys. J. 120, 924–935 (2021).

26. Barstow, B., Ando, N., Kim, C. U. & Gruner, S. M. Alteration of citrine structure by hydrostatic pressure explains the accompanying spectral shift. Proc. Natl. Acad. Sci. U. S. A. 105, 13362–13366 (2008).

27. Kundrot, C. E. & Richards, F. M. Crystal structure of hen egg-white lysozyme at a hydrostatic pressure of 1000 atmospheres. J. Mol. Biol. 193, 157–170 (1987).

28. Urayama, P., Phillips, G. N., Jr & Gruner, S. M. Probing substates in sperm whale myoglobin using high-pressure crystallography. Structure 10, 51–60 (2002).

29. Collins, M. D., Hummer, G., Quillin, M. L., Matthews, B. W. & Gruner, S. M. Cooperative water filling of a nonpolar protein cavity observed by high-pressure crystallography and simulation. Proc. Natl. Acad. Sci. U. S. A. 102, 16668–16671 (2005).

30. Collins, M. D., Quillin, M. L., Hummer, G., Matthews, B. W. & Gruner, S. M. Structural rigidity of a large cavity-containing protein revealed by high-pressure crystallography. J. Mol. Biol. 367, 752–763 (2007).

31. Yamada, H., Nagae, T. & Watanabe, N. High-pressure protein crystallography of hen egg-white lysozyme. Acta Crystallogr. D Biol. Crystallogr. 71, 742–753 (2015).

32. Girard, E. et al. Equilibria between conformational states of the Ras oncogene protein revealed by high pressure crystallography. Chem. Sci. 13, 2001–2010 (2022).

33. Prangé, T. et al. Comparative study of the effects of high hydrostatic pressure per se and high argon pressure on urate oxidase ligand stabilization. Acta Crystallogr D Struct Biol 78, 162–173 (2022).

34. Kurup, P., Zhang, Y., Venkitaramani, D. V., Xu, J. & Lombroso, P. J. The role of STEP in Alzheimer’s disease. Channels 4, 347–350 (2010).

35. Goebel-Goody, S. M. et al. Genetic manipulation of STEP reverses behavioral abnormalities in a fragile X syndrome mouse model. Genes Brain Behav. 11, 586–600 (2012).

36. Kurup, P. K. et al. STEP61 is a substrate of the E3 ligase parkin and is upregulated in Parkinson’s disease. Proceedings of the National Academy of Sciences 112, 1202–1207 (2015).

37. Berman, H. M. et al. The Protein Data Bank. Nucleic Acids Res. 28, 235–242 (2000).

38. Barr, A. J. et al. Large-scale structural analysis of the classical human protein tyrosine phosphatome. Cell 136, 352–363 (2009).

39. Tautermann, C. S. et al. Allosteric Activation of Striatal-Enriched Protein Tyrosine Phosphatase (STEP, PTPN5) by a Fragment-like Molecule. J. Med. Chem. 62, 306–316 (2019).

40. Ginn, H. M. Torsion angles to map and visualize the conformational space of a protein. Protein Sci. e4608 (2023).

41. Eswaran, J. et al. Crystal structures and inhibitor identification for PTPN5, PTPRR and PTPN7: a family of human MAPK-specific protein tyrosine phosphatases. Biochem. J 395, 483–491 (2006).

42. Witten, M. R. et al. X-ray Characterization and Structure-Based Optimization of Striatal-Enriched Protein Tyrosine Phosphatase Inhibitors. J. Med. Chem. 60, 9299–9319 (2017).

43. Kim, C. U., Kapfer, R. & Gruner, S. M. High-pressure cooling of protein crystals without cryoprotectants. Acta Crystallogr. D Biol. Crystallogr. 61, 881–890 (2005).

44. Chen, C. R. & Makhatadze, G. I. ProteinVolume: calculating molecular van der Waals and void volumes in proteins. BMC Bioinformatics 16, 101 (2015).

45. Tyka, M. D. et al. Alternate states of proteins revealed by detailed energy landscape mapping. J. Mol. Biol. 405, 607–618 (2011).

46. Tautz, L., Critton, D. A. & Grotegut, S. Protein tyrosine phosphatases: structure, function, and implication in human disease. Methods Mol. Biol. 1053, 179–221 (2013).

47. Lovell, S. C. et al. Structure validation by Calpha geometry: phi,psi and Cbeta deviation. Proteins 50, 437–450 (2003).

48. Lovell, S. C., Word, J. M., Richardson, J. S. & Richardson, D. C. The penultimate rotamer library. Proteins 40, 389–408 (2000).

49. Joosten, R. P., Long, F., Murshudov, G. N. & Perrakis, A. The PDB_REDO server for macromolecular structure model optimization. IUCrJ 1, 213–220 (2014).

50. Fourme, R. et al. High-pressure protein crystallography (HPPX): instrumentation, methodology and results on lysozyme crystals. J. Synchrotron Radiat. 8, 1149–1156 (2001).

51. Ascone, I., Savino, C., Kahn, R. & Fourme, R. Flexibility of the Cu,Zn superoxide dismutase structure investigated at 0.57 GPa. Acta Crystallogr. D Biol. Crystallogr. 66, 654–663 (2010).

52. Williamson, M. P., Akasaka, K. & Refaee, M. The solution structure of bovine pancreatic trypsin inhibitor at high pressure. Protein Sci. 12, 1971–1979 (2003).

53. Critton, D. A., Tautz, L. & Page, R. Visualizing active-site dynamics in single crystals of HePTP: opening of the WPD loop involves coordinated movement of the E loop. J. Mol. Biol. 405, 619–629 (2011).

54. Yeh, C. Y. et al. A conserved local structural motif controls the kinetics of PTP1B catalysis. bioRxiv 2023.02.28.529746 (2023) doi:10.1101/2023.02.28.529746.

55. Whittier, S. K., Hengge, A. C. & Loria, J. P. Conformational motions regulate phosphoryl transfer in related protein tyrosine phosphatases. Science 341, 899–903 (2013).

56. Braithwaite, S. P., Paul, S., Nairn, A. C. & Lombroso, P. J. Synaptic plasticity: one STEP at a time. Trends Neurosci. 29, 452–458 (2006).

57. Chen, Y.-N. P. et al. Allosteric inhibition of SHP2 phosphatase inhibits cancers driven by receptor tyrosine kinases. Nature 535, 148–152 (2016).

58. Yu, X. et al. Structure, inhibitor, and regulatory mechanism of Lyp, a lymphoid-specific tyrosine phosphatase implicated in autoimmune diseases. Proc. Natl. Acad. Sci. U. S. A. 104, 19767–19772 (2007).

59. Winter, G. et al. DIALS as a toolkit. Protein Sci. 31, 232–250 (2022).

60. Karplus, P. A. & Diederichs, K. Linking crystallographic model and data quality. Science 336, 1030–1033 (2012).

61. Wojdyr, M., Keegan, R., Winter, G. & Ashton, A. DIMPLE - a pipeline for the rapid generation of difference maps from protein crystals with putatively bound ligands. Acta Crystallogr. A 69, 299–299 (2013).

62. Murshudov, G. N., Vagin, A. A. & Dodson, E. J. Refinement of macromolecular structures by the maximum-likelihood method. Acta Crystallogr. D Biol. Crystallogr. 53, 240–255 (1997).

63. Afonine, P. V. et al. Towards automated crystallographic structure refinement with phenix.refine. Acta Crystallogr. D Biol. Crystallogr. 68, 352–367 (2012).

64. Emsley, P. & Cowtan, K. Coot: model-building tools for molecular graphics. Acta Crystallogr. D Biol. Crystallogr. 60, 2126–2132 (2004).

65. Liebschner, D. et al. Macromolecular structure determination using X-rays, neutrons and electrons: recent developments in Phenix. Acta Crystallogr D Struct Biol 75, 861–877 (2019).

66. Williams, C. J. et al. MolProbity: More and better reference data for improved all-atom structure validation. Protein Sci. 27, 293–315 (2018).

67. Humphrey, W., Dalke, A. & Schulten, K. VMD: visual molecular dynamics. J. Mol. Graph. 14, 33–8, 27–8 (1996).

68. Hintze, B. J., Lewis, S. M., Richardson, J. S. & Richardson, D. C. Molprobity’s ultimate rotamer-library distributions for model validation. Proteins 84, 1177–1189 (2016).

69. Pedersen, A. K., Peters G G. Ü. H., Møller, K. B., Iversen, L. F. & Kastrup, J. S. Water-molecule network and active-site flexibility of apo protein tyrosine phosphatase 1B. Acta Crystallogr. D Biol. Crystallogr. 60, 1527–1534 (2004).

