## Supplementary Figures for "Pushed to extremes: distinct effects of high temperature vs. pressure on the structure of an atypical phosphatase"

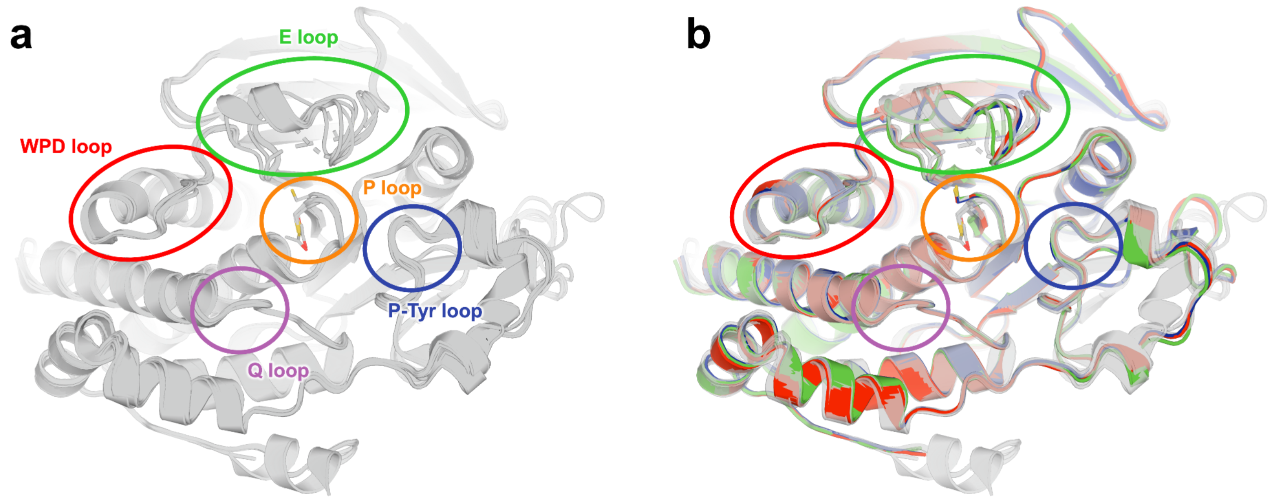


**Supplementary Figure 1: Superposition of our new STEP structures and all previous STEP structures.**

a) All previous human and mouse structures overlaid: PDB IDs 2bv5, 2bij, 2cjz, 5ovr, 5ovx, 5ow1, 6h8r, and 6h8s (gray).

b) Our LoTP (blue), HiT (red), and HiP (green) structures are shown overlaid with all previous human and mouse STEP structures.

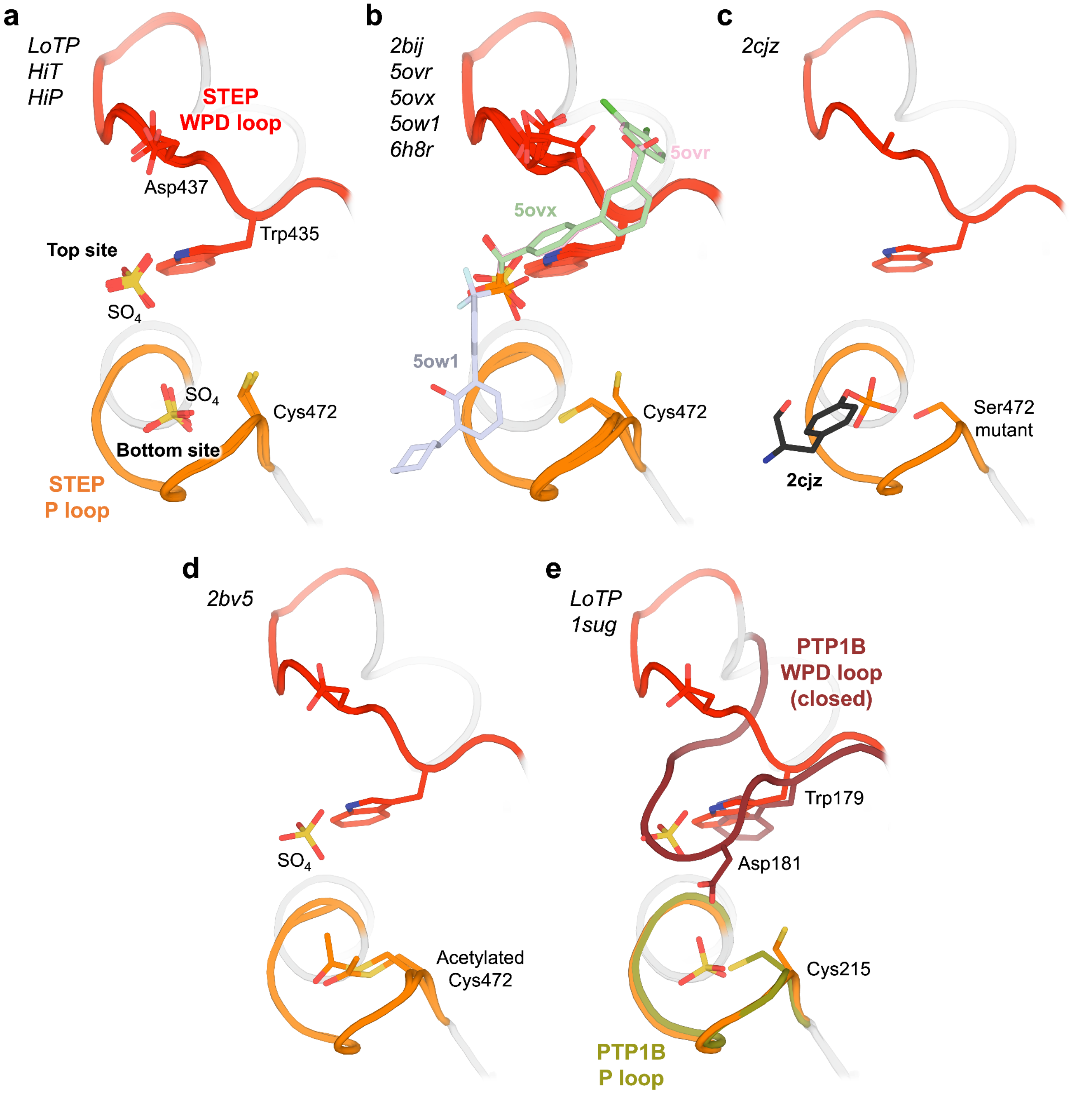


**Supplementary Figure 2: Previous STEP structures did not have two sulfates in the active site.**

a) Sulfates in our LoTP, HiT, and HiP structures in both the “top” and “bottom” sites. Key residues in the WPD loop and the P loop are labeled.

b) A sulfate or analogous group from a competitive inhibitor only in the top site, in PDB ID 2bij, 5ovr, 5ovx, 5ow1, and 6h8r.

c) A phosphate group in a pTyr substrate in PDB ID only in the bottom site, in PDB ID 2cjz.

d) A sulfate in the top site and an acetylated catalytic Cys472 in the bottom site, in PDB ID 2bv5.

e) Superposition of the PTP1B closed state from PDB ID 1sug [69] with our STEP LoTP structure, illustrating that the closed WPD loop of PTP1B aligns well with the top sulfate in STEP. Key residues from PTP1B in the WPD loop and P loop are labeled.

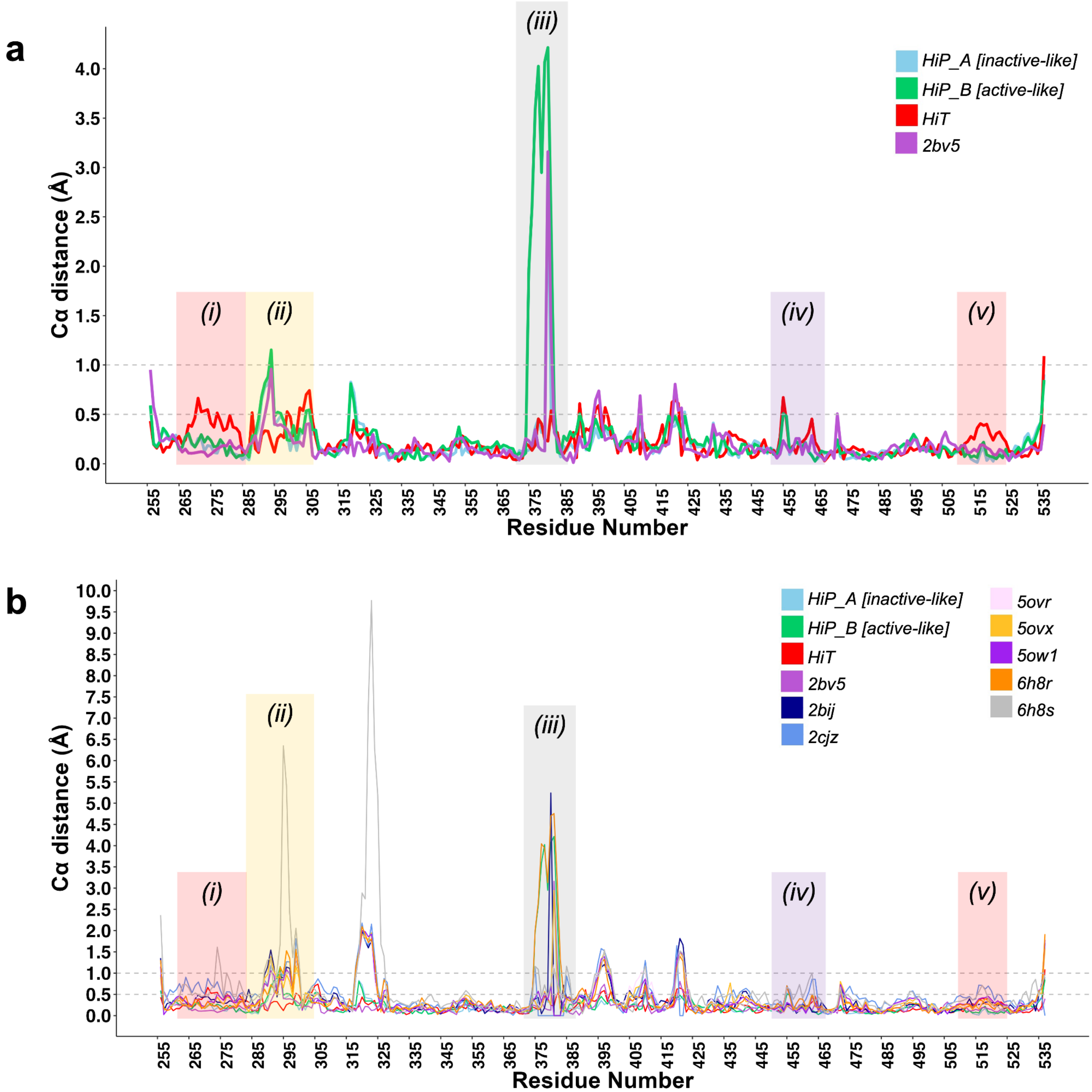


**Supplementary Figure 3: Global backbone displacements due to high temperature vs. pressure, relative to previous STEP structures.**

a) Cα distances for the HiT and HiP structures relative to the reference LoTP structure vs. amino acid sequence, including the only previous STEP structure in the same crystal form as our structures (PDB ID 2bv5).

b) Same as (a), but also including all previous human and mouse STEP structures thathave a different crystal form (PDB ID 2bij, 2cjz, 5ovr, 5ovx, 5ow1, 6h8r, 6h8s).

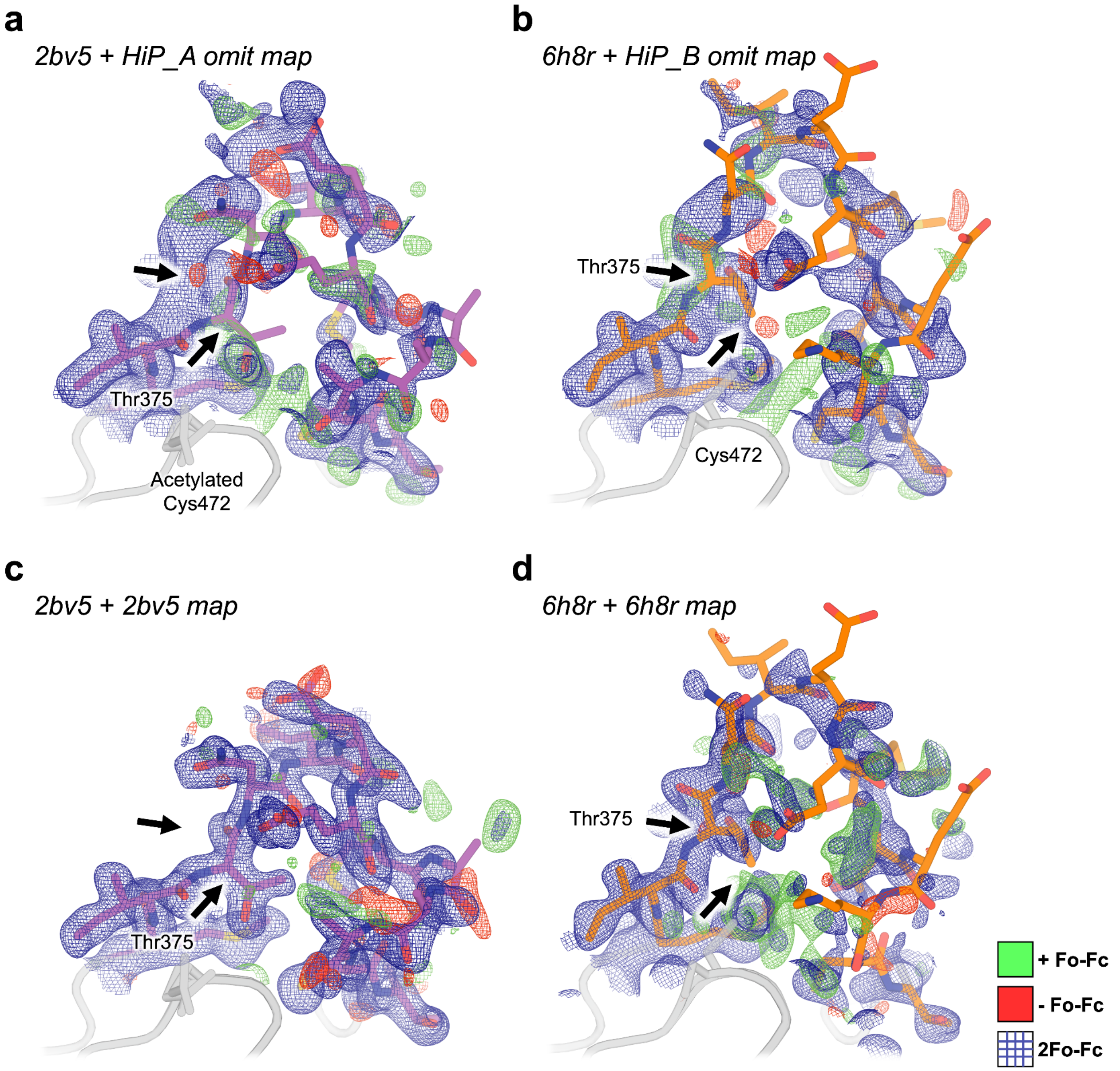


**Supplementary Figure 4: The E loop at high pressure samples two distinct previously observed states.**

a) PDB 2bv5 model with our HiP 2Fo-Fc (contoured at 1 σ) and Fo-Fc (contoured at ±3 σ) maps omitting alternate conformation A.
b) PDB 6h8r model with our HiP 2Fo-Fc and Fo-Fc maps omitting alternate conformation B.
c) 2bv5 model with 2bv5 2Fo-Fc and Fo-Fc maps.
d) 6h8r model with 6h8r 2Fo-Fc and Fo-Fc maps.

Arrows highlight the locations of the dual-conformation Thr375 in our HiP model, for reference.

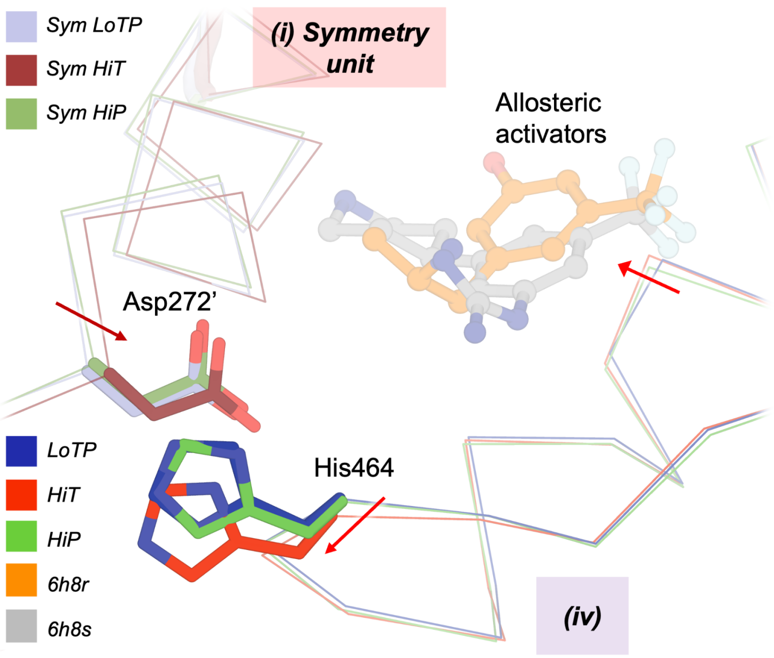


**Supplementary Figure 5: Backbone displacements at the allosteric activator site due to high temperature.**
Zoom-in of the area including region (iv) (residues 454-467, S loop) from **Fig. 3a**. The small-molecule allosteric activators from PDB ID 6h8r (orange) and 6h8s (gray) are shown for context, as well as the symmetry-related Asp272’ from region (i) from **Fig. 3a**.

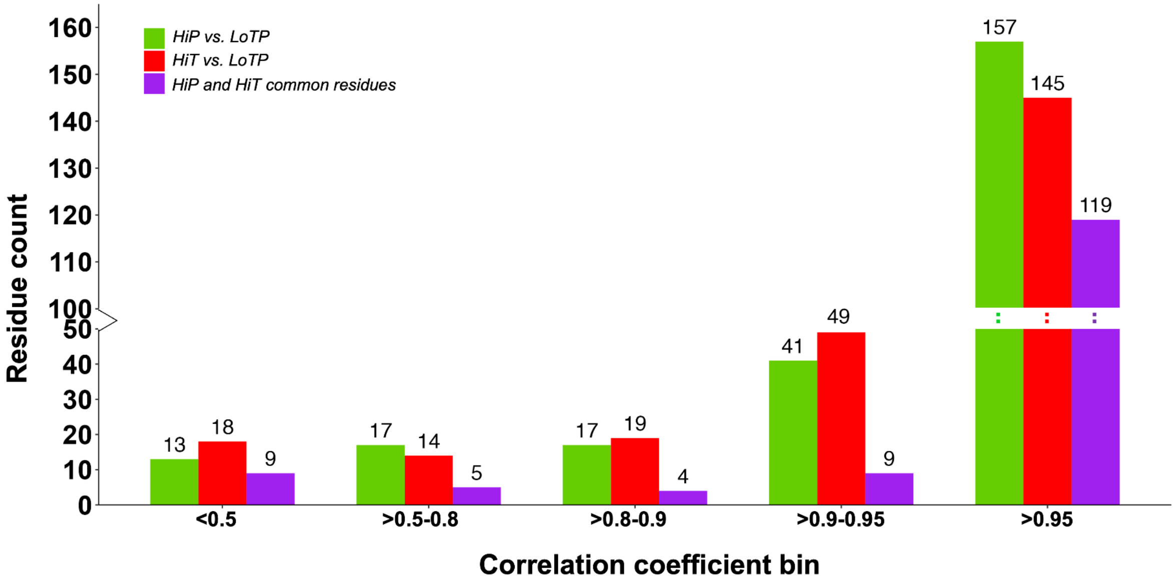


**Supplementary Figure 6: Ringer analysis points to widespread but distinct effects of high temperature vs. pressure on side-chain conformations.**
For each residue in STEP (excluding the flexible E loop), Ringer curves were calculated for HiT, HiP, and LoTP, and Pearson correlation coefficient (CC) was calculated relative to LoTP. Shown are the number of residues with given Ringer CC values for HiT, for HiP, and for both HiT + HiP.

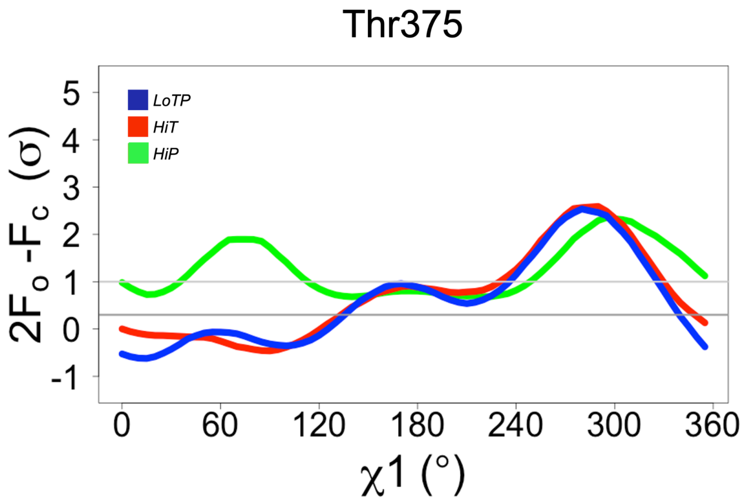


**Supplementary Figure 7: Differences in Ringer curves at high pressure due to distinct backbone positioning.**

Thr375 has similar χ1 peaks for LoTP and HiT, but different χ1 peaks for HiP. (Note that Thr generally has two χ1 peaks even with a single conformation because it is a β-branched side chain with two different γ atoms.) This is associated with backbone shifts starting at Thr375 at the beginning of the E loop that occur only at HiP. LoTP in blue, HiT in red, HiP in green, as in **Fig. 5**.
